# TuMV infection alters miR168/AGO1 and miR403/AGO2 systems regulation in Arabidopsis

**DOI:** 10.1101/2021.02.17.431672

**Authors:** Carlos Augusto Manacorda, Sabrina Tasselli, María Rosa Marano, Sebastian Asurmendi

## Abstract

Plant Argonaute proteins, chiefly AGO1 and 2, restrict viral infections. AGO1/2 also participate in developmental processes and are tightly regulated by microRNAs. Under viral infections, the regulatory loop comprising miR168/AGO1 is well studied, but much less so the miR403/AGO2 system. We studied both regulatory systems in TuMV-infected Arabidopsis plants. TuMV downregulated miRNAs precursor molecules, but mature miRNAs overaccumulated, without evidence of transcriptional alteration. AGO1 protein remained at basal levels whereas AGO2 overaccumulated. These results are in line with previous reports studying abiotic and biotic impact on microRNA biogenesis and AGO-dependent antiviral defense, expanding our knowledge of the miR403/AGO2 regulatory system.

## 1. Introduction

Potyvirus is the largest plant virus genus, causing significant losses in a wide range of crops (Revers and García, 2015). Turnip Mosaic Virus (TuMV) is one of the best studied potyviruses. It can infect not only the model plant Arabidopsis but also dozens of species of economic importance including commercial brassicas (Walsh and Jenner, 2002). Plant viral infections trigger a strong response of the host RNA silencing machinery (Yang and Li, 2018). In Arabidopsis, ten ARGONAUTE (AGO) proteins exist, which associate either with microRNAs (miRNAs), thus regulating the translation of endogenous messenger RNAs, or with short interfering RNAs (siRNAs), a feature that is paramount in the plant antiviral RNA silencing defense, where AGO proteins utilize virus-derived siRNAs to target and cut viral RNA molecules (Vaucheret, 2008; Fang and Qi, 2015). Among these AGO proteins, AGO1/2/10 have been shown to play a key role in Arabidopsis during TuMV infection, with AGO2 playing a major role in leaves (Garcia-ruiz et al., 2015). The protein AGO1, which is itself regulated at the mRNA level by miR168 (Vazquez et al., 2004) has a well-established role in normal development but also in viral siRNA loading and silencing (Vance and Vaucheret, 2001). The homeostasis of AGO1 is partially dependent on the post-transcriptional regulatory loop miR168/AGO1 (Vaucheret et al., 2006). This homeostasis is perturbed by several unrelated viruses affecting plants that have convergently evolved a molecular counter-attack based on inactivation of the AGO1–based plant defense by dramatically increasing miR168 accumulation, therefore limiting the amount of AGO1 protein and compromising its antiviral function (Várallyay and Havelda, 2013; Várallyay et al., 2010). Mature miR168 originates from two precursor molecules derived from different loci, miR168a and b. It has been shown that several viral infections can induce miR168 accumulation by means of transcriptional activation of miR168a gene (Várallyay and Havelda, 2013). Interestingly, miR168 and AGO1 are positively regulated by ABA (Li et al., 2012), which also affects the maturation and stability of miRNAs (Alazem et al., 2014; Alazem and Lin, 2017). TuMV enhances mature miR168 overaccumulation (Garcia-ruiz et al., 2015) and also ABA and SA levels (Poque et al., 2018; Manacorda et al., 2021). Both hormones have positively correlated functions in antiviral activity, including silencing activation (Alazem et al., 2019; Pasin et al., 2020; Alazem and Lin, 2017; Yang and Li, 2018). Also ABA and particularly, SA, were found to induce AGO2 in Arabidopsis and Nicotiana benthamiana (Diao et al., 2019; Alazem et al., 2017; Alazem et al., 2019). In leaves, an emerging consensus places AGO2 as a main player in antiviral silencing (Carbonell and Carrington, 2015; Yang and Li, 2018) including TuMV (Carbonell et al., 2012; Ludman et al., 2017; Garcia-ruiz et al., 2015), mainly due to the effective AGO1 repression by viruses (Harvey et al., 2011; Várallyay and Havelda, 2013). Under normal conditions, AGO2 is in turn post-transcriptionally regulated by miR403 loaded into AGO1 (Allen et al., 2005). However, only recently the regulation of the miR403/AGO2 system has been investigated during viral infections (Xu et al., 2016; Pertermann et al., 2018; Diao et al., 2019). However, the simultaneous study of both miR168/AGO1 and miR403/AGO2 regulatory systems during a viral infection is still lacking. Here, we investigated the impact of Arabidopsis infection with TuMV on the miR168/AGO1 and miR403/AGO2 antiviral silencing systems. We used a time-course methodology previously developed that had revealed useful to study transcriptional, biochemical and morphological changes upon TuMV infection (Manacorda et al., 2013), from early (4 DPI) to advanced (16 DPI) stages of viral infection, in systemically infected rosette leaves. We also evaluated the effect of TuMV infection on the accumulation of AGO1 and AGO2 proteins.

## 2. Materials and Methods

### 2.1. Plant growing conditions and material

*Arabidopsis thaliana* Col-0 and pmiR168a:GUS transgenic plants were grown as described in (Manacorda et al., 2021) for both long and short days conditions. Leaves number 8, 11 and 13 were sampled as these leaves constituted the vascular sympodia connecting leaf #3 (the inoculated one) (Kang et al., 2003; Heyer et al., 2018) and are likely to receive the full systemic impact of TuMV infection (Manacorda and Asurmendi, 2018; Manacorda et al., 2013). For short days conditions, systemic leaves were pooled from different plants to obtain sufficient plant material to be jointly grinded and separated in aliquots for subsequent use in qPCR, Western blot and Northern blot experiments. Two independent experiments were performed showing similar results. Biological replicates consisted of 5 ≥ N ≥ 3 pools per experiment and treatment and each pool consisted of 5 ≥ N ≥ 3 plants.

### 2.2. Virus infection assays

Viral infections were performed similarly as described in (Manacorda et al., 2021). Only leaves #3 were inoculated either with virus or buffer (mock-inoculations). TuMV-UK1 strain (accession number AF169561) (Sánchez et al., 1998) and JPN1 strain (accession number KM094174) (Sánchez et al., 2015) were used. Effectiveness of viral infections was visually confirmed based on long-term symptoms development as in (Manacorda et al., 2021). Only plants developing characteristic TuMV symptoms were used (**Supplementary Figure SF1**).

### 2.3. qPCR assays

Oligonucleotide primer sets for real-time (quantitative) PCR were designed using Vector NTI Advance 9 software (Life Technologies, Carlsbad, CA, U.S.A.). They are listed in **Supplementary Table TS3**. The design of microRNAs primers for cDNA synthesis and qPCR assays were adapted from (Chen et al., 2005; Pant et al., 2008; Benes and Castoldi, 2010). Primer design for pre-miRNAs were done taking into account the processing steps (all of them in a base-to-loop fashion (Bologna et al., 2013)) that lead from pri- to pre-miRNA to encompass the mature microRNA, in a sense similar to what have been proposed to qPCR-amplify animal pre-miRNAs (Schmittgen et al., 2008). Details on the minimum information for publication of quantitative real-time PCR experiments requirements are listed in **Supplementary Tables ST1 and ST2**.

### 2.4. Northern blot analyses

Northern blots for miRNA analysis were performed as previously described (Dalmay, 2011; Giacomelli *et al*., 2012). The signals were detected by chemiluminescence (ECL, Thermo Scientific, Petaluma, CA, USA) and exposure to autoradiograph films (BioMax MS/Screen, Kodak, NY, USA). U6 was used as the internal standard control. Probe sequences are listed in **Supplementary Table ST3**.

### 2.5. Immunoblot analyses

Proteins were isolated from 300 mg of ground tissue with 300 μL of extraction buffer (50 mM Tris, pH 7.5; 150 mM NaCl; 1 mM EDTA; 10% (v/v) glycerol; 1 mM dithiothreitol (DTT); 1 mM Pefabloc (Roche) and one tablet of complete protease inhibitor cocktail (Roche)) vortexing for 10 s. Tissue debris was eliminated by centrifugation at 12 000 g at 4°C for 20 m. Fifty microliters of crude protein extract per sample were resolved on an 8% polyacrylamide gel and transferred to a polyvinylidene difluoride membrane (PVDF-Immun-Blot®, Bio-Rad, Hercules, CA, USA) as described previously (Qi and Mi, 2010). Immunoblotting was performed using a polyclonal antibody with reactivity against Arabidopsis AGO1 (1:10000 dilution) or AGO2 (1:250 dilution) (Agrisera, Vännäs, Sweden) and subsequently detected by a horseradish peroxidase (HRP)-conjugated antirabbit IgG (Agrisera, Vännäs, Sweden, 1:20000 dilution), and chemiluminescence (ECL, Thermo Scientific, Petaluma, CA, USA) according to manufacturer’s indications. For AGO1 and AGO2 proteins the results of two similar experiments were combined and jointly analyzed. For AGO1* and AGO2* forms only one experiment detected bands in mock-inoculated controls and was then suitable to be statistically analyzed. 8 ≥ N ≥ 3 per group.

### 2.6. Image analyses

Image analyses of Northern and Western blot hybridized membranes were performed using ImageJ (Schneider *et al*., 2012) using the Gels Submenu as suggested in https://imagej.nih.gov/ij/docs/menus/analyze.html#gels after background subtraction (https://imagej.nih.gov/ij/docs/examples/dot-blot/index.html). MiRNAs and AGO proteins signals were relativized to U6 and RuBisCo (from the protein Ponceau staining) respectively and then normalized to their mock-inoculated respective controls.

### 2.7. Hormone treatment

SA treatment was performed as described in (Manacorda *et al*., 2013).

### 2.8. Statistical analysis

For all statistical analysis, significance was set as: NS (Not Significative statistical differences) = P > 0.05, * = P ≤ 0.05, ** = P ≤ 0.01, and *** = P ≤ 0.001. For data presenting nested (hierarchical) structure, linear mixed effects models (*lme*) were implemented in R (RStudio Team, 2020; RCoreTeam, 2020) using the *nlme* package (Pinheiro *et al*., 2020), used as suggested in (Pinheiro and Bates, 2000). For qPCR analysis, LinRegPCR program (Ramakers *et al*., 2003) and the normalization method of (Pfaffl *et al*., 2002) were used. Relative expression ratios and statistical analysis were done using fgStatistics software interface (Di Rienzo, 2009).

## 3. Results

### 3.1. TuMV infection alters miR168 biogenesis and the miR168/AGO1 system regulation

To examine the effect of TuMV (UK1 strain) infection on miR168/AGO1 system, we inoculated Arabidopsis plants growing in long days (LD) conditions and collected systemic aerial tissue at several days post-inoculation (DPI) (**Supplementary Tables ST1 and ST2**). Mature miR168 was highly overaccumulated under infection (**Fig. 1a**). However, when levels of precursor molecules that contribute to mature miR168 accumulation were analyzed, a downregulation was detected under TuMV infection (**Fig. 1b,c**). We tested whether these diminished levels of precursor molecules were linked to lower transcriptional levels at the promoter level. As miR168a is ubiquitously expressed and coexpressed with AGO1 whereas miR168b expression is restricted to the shoot apical meristem (Vaucheret, 2009), pmiR168a:GUS transgenic plants (Vaucheret et al., 2006) were surveyed for transcriptional activity under TuMV infection. Given the high stability of GUS protein, GUS mRNA levels were measured instead (Dolata et al., 2016; Várallyay and Havelda, 2013). **Fig. 1d** shows little or no repression of GUS mRNA after infection even though endogenous pre-miR168a was also significantly down-regulated in these pmiR168a:GUS transgenic plants, suggesting that TuMV has no transcriptional effect upon pmiR168a. Despite mature miR168 overaccumulation during TuMV infection, AGO1 mRNA did not decrease during the entire period analyzed but on the contrary, an overaccumulation was detected at 10 DPI (**Fig. 1e**). Another TuMV strain, JPN1, was reported to induce different phenotypical and molecular responses (Sánchez et al., 2015; Manacorda et al., 2013) either in their nature or in timing of the response, but has also shown some similar physiological outcomes in Arabidopsis (Manacorda et al., 2021). We then tested how the miR168/AGO1 system behaved when the JPN1 strain of TuMV, is used (**Fig. SF2**). Results showed that essentially the same molecular response is triggered by Arabidopsis regarding miR168/AGO1 system. Interestingly, JPN1 strain also down-regulated pre-miR168a despite the absence of any transcriptional effect detected on GUS reporter gene in pmiR168a:GUS transgenic plants (**Fig. SF2d**). This further suggested that pre-miR168a down-regulation by TuMV arises from post-transcriptional events. Overall, these results showed that TuMV alters miR168 biogenesis at a post-transcriptional stage and that mature miR168 overaccumulation fails to downregulate AGO1 mRNA below basal levels. Given that both TuMV strains induced similar molecular responses regarding miR168/AGO1 system, we selected the UK1 strain to continue our work.

**Fig. 1.**
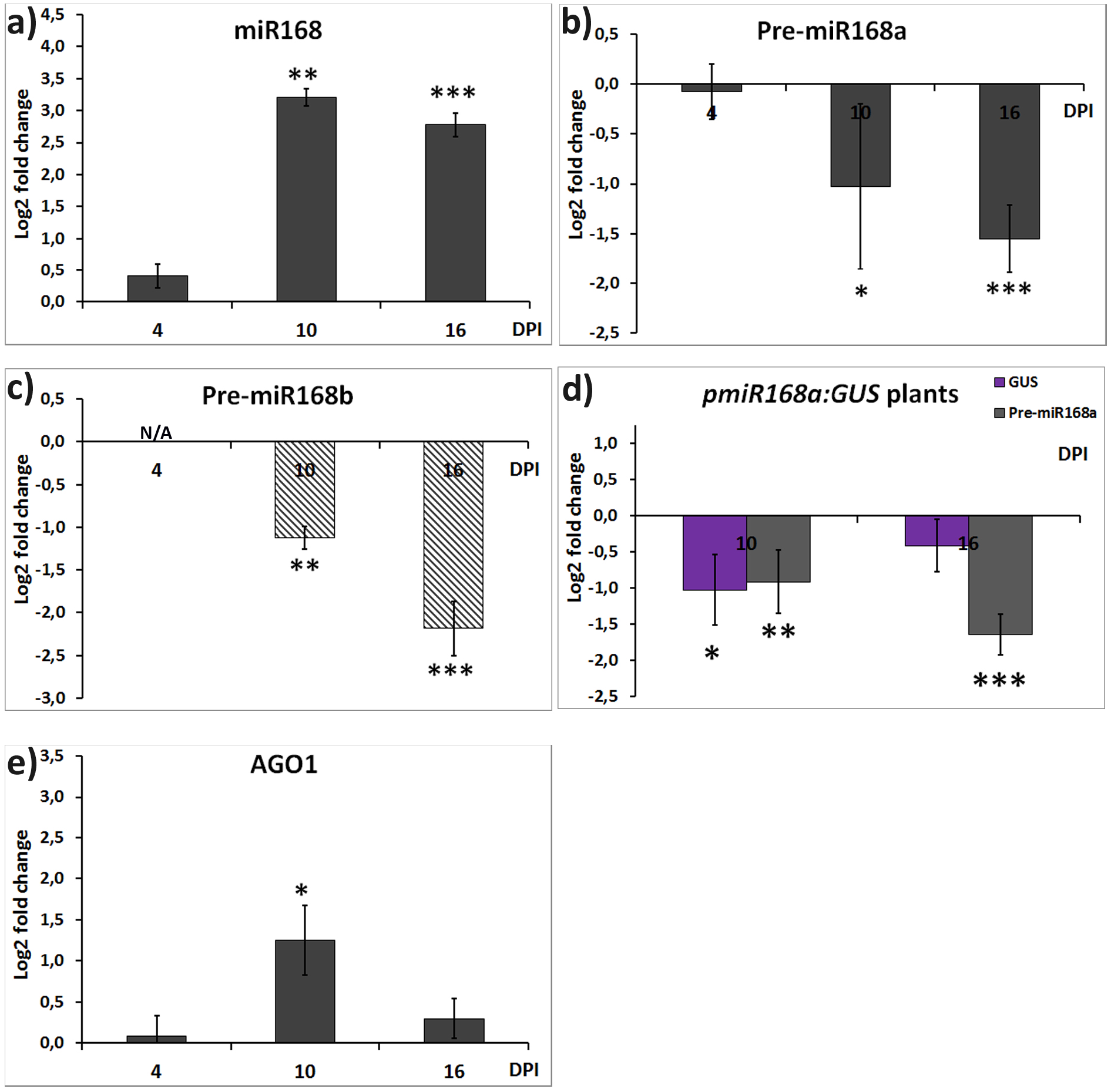
TuMV alters miR168 accumulation and biogenesis and miR168/AGO1 system regulation. Time-course qPCR analyses measuring relative accumulation of **a)** mature miR168, **b-c)** miR168a/b precursor forms, **d)** GUS mRNA and endogenous pre-miR168a in *pmiR168a:GUS* transgenic plants and **e)** AGO1 mRNA, in TuMV-infected *versus* control (mock-inoculated) plants. **A-c, e)** Col-0 WT plants; **d)** *pmiR168a:GUS* transgenic plants. Bars = SE. 6 ≥ N ≥ 3 per group. N/A = Not Analyzed.

### 3.2. TuMV infection alters miR403 biogenesis and the miR403/AGO2 system regulation

TuMV induced the overaccumulation of mature miR403 (**Fig. 2a**) but pre-miR403 was downregulated by TuMV (**Fig. 2b**). AGO2 mRNA was overaccumulated from early stages of infection throughout the entire time-frame analyzed (**Fig. 2c**). The hormonal balance has been found to be important for AGO genes expression (Alazem *et al*., 2019). Previous studies have found a transcriptomic pattern of expression that is coherent with the overaccumulation of SA triggered by TuMV (Sánchez *et al*., 2015; Manacorda *et al*., 2021) even at early stages of infection (Manacorda *et al*., 2013). We therefore investigated the response of AGO1 and AGO2 gene expression after SA foliar spray application. A transient high induction was observed for AGO2, whereas AGO1 remained at basal levels (**Fig. 3)**. Summarizing, **Fig. 2 and Fig. 3** showed that TuMV alters miR403 biogenesis and that mature miR403 overaccumulation occurs concomitantly with AGO2 mRNA overaccumulation, a result that is coherent with TuMV being able to induce high levels of SA. Since our previous work (Manacorda *et al*., 2021) showed that physiological responses to TuMV are conserved between different growing conditions, we repeated molecular analyses for miR168/AGO1 and miR403/AGO1 regulatory systems in short days (SD) conditions to assess the robustness of the investigated phenomena. Northern blot and qPCR analyses showed that both regulatory systems underwent essentially the same changes following TuMV infection in SD (**Fig. 4**) compared with results in LD (**Fig. 1 and Fig. 2**). Therefore, TuMV deregulated miR168/AGO1 and miR403/AGO2 systems in both LD and SD conditions.

**Fig. 2.**
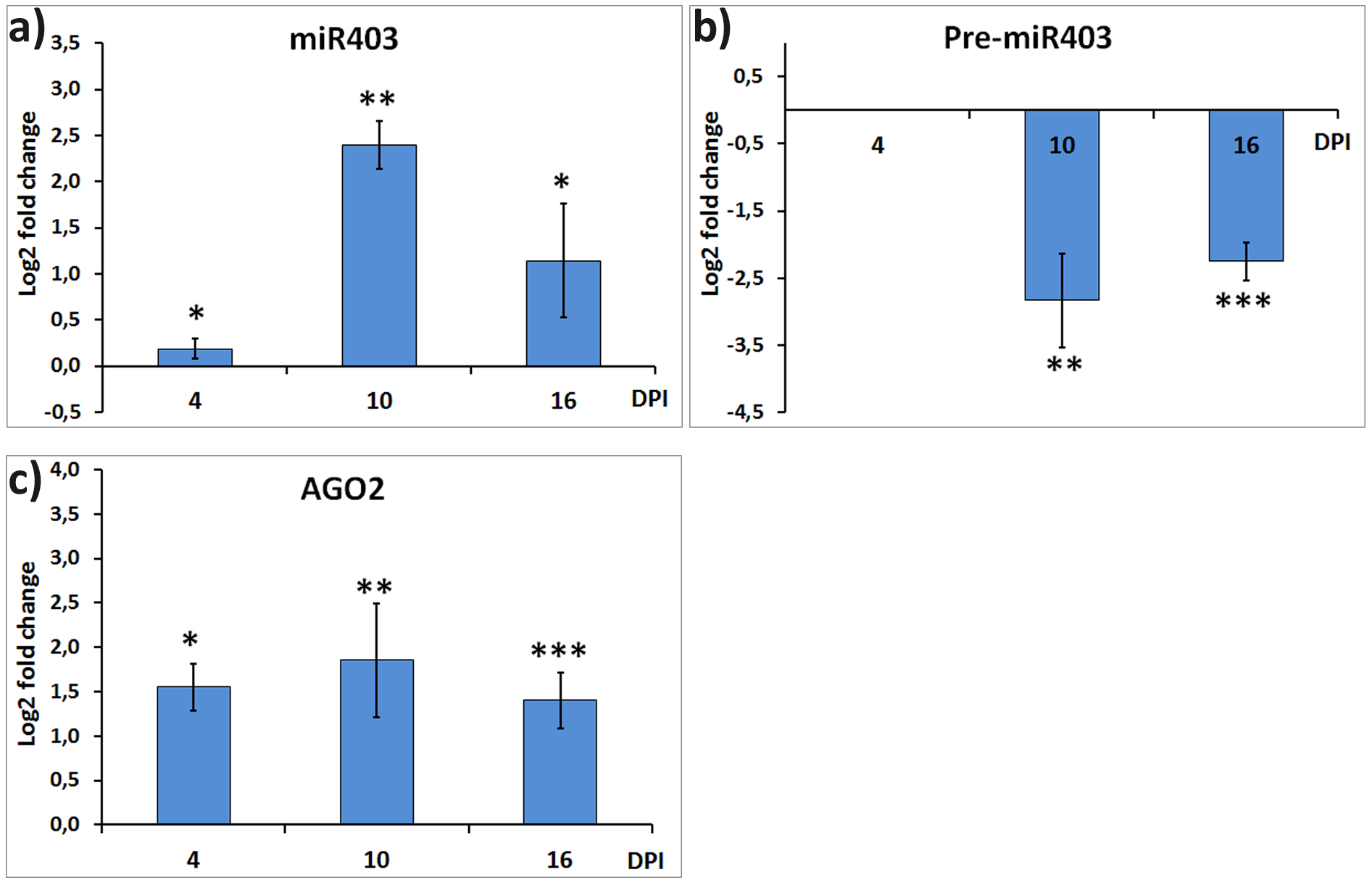
TuMV alters miR403 accumulation and biogenesis and miR403/AGO2 system regulation. QPCR analyses measuring relative accumulation of **a)** mature miR403, **b)** pre-miR403, **c)** AGO2 mRNA, in TuMV-infected *versus* control mock-inoculated) Col-0 WT plants. Bars = SE. 7 ≥ N ≥ 3 per group. N/A = Not Analyzed.

**Fig. 3:**
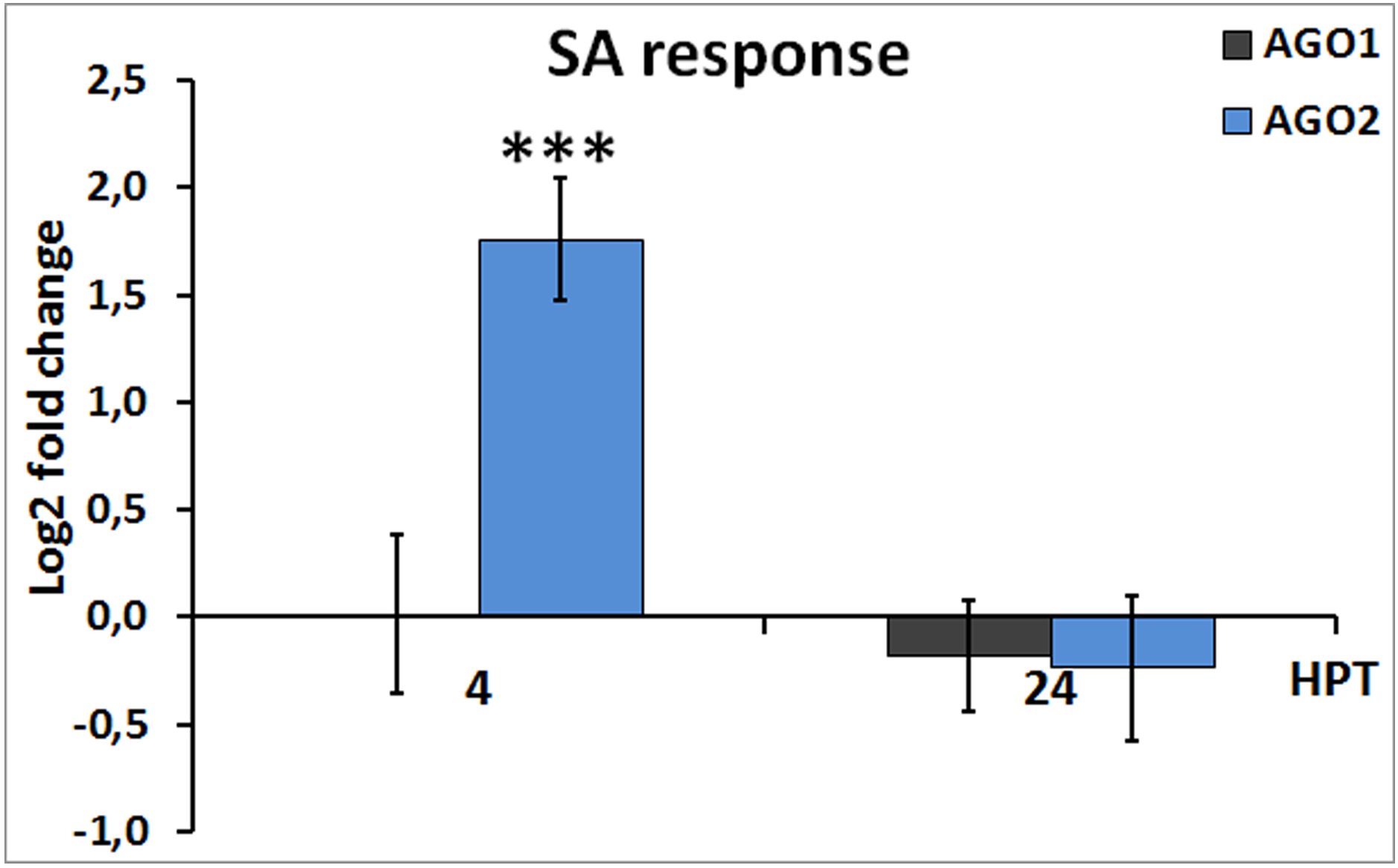
AGO2, but not AGO1, is SA-responsive. QPCR analyses measuring relative accumulation of AGO1 and AGO2 mRNAs in SA-sprayed *versus* control (water-sprayed) plants. Bars = SE. N = 5 per group.

**Fig. 4:**
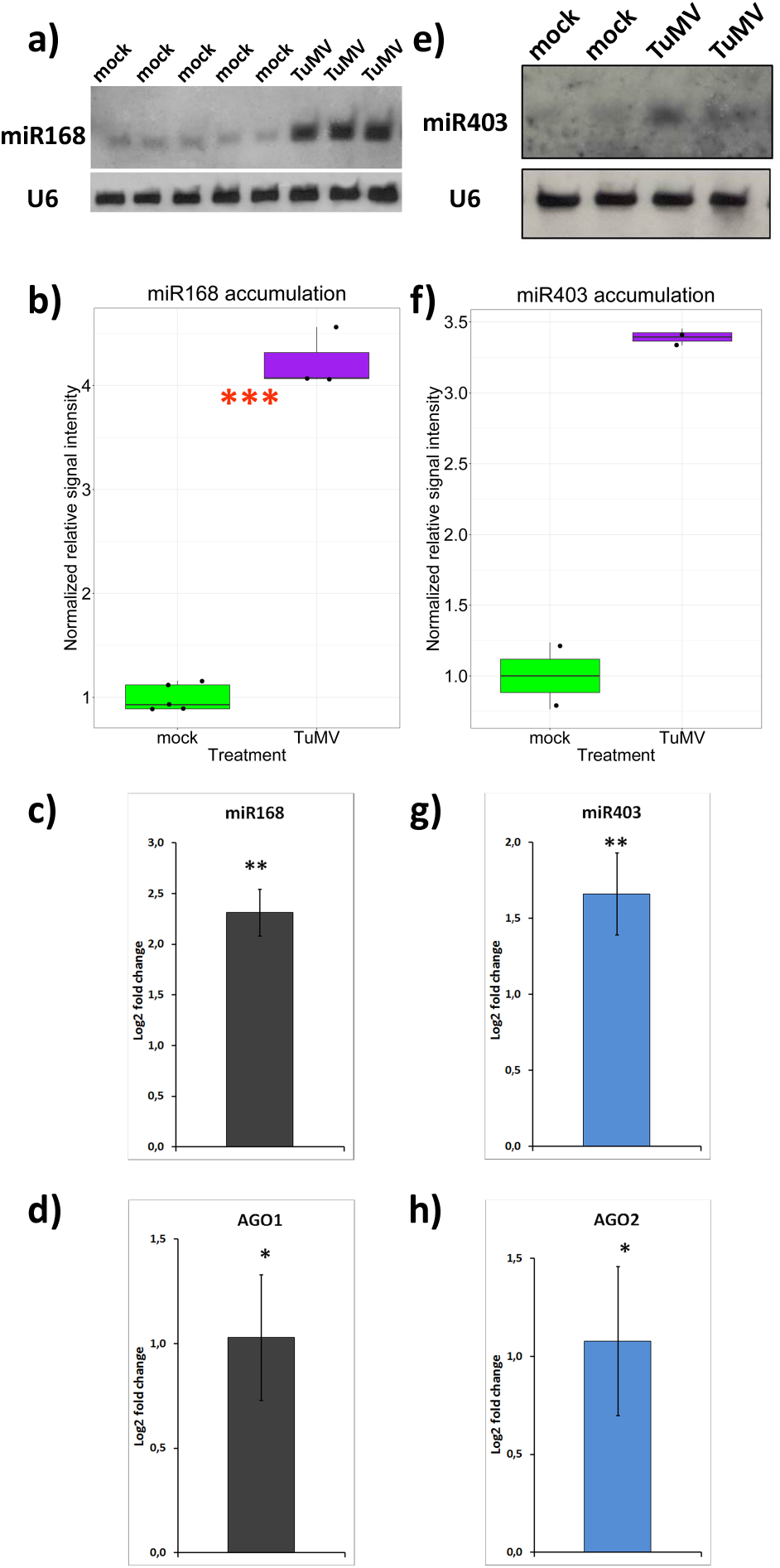
Mir168/AGO1 and miR403/AGO2 systems are also altered by TuMV in short days (SD) conditions. **a-d)** miR168/AGO1 system and **e-h)** miR403/AGO2 system. **(a,b)** and **(e,f):** Northern blot results with corresponding box-and-jitter plots analyses for relative accumulation of miR168 and miR403, respectively, under TuMV infection. 5 ≥ N ≥ 2 per group. **(c,d)** and **(g,h):** qPCR analyses of **c)** miR168, **d)** AGO1, **g)** miR403 and **h)** AGO2. Bars = SE. 5 ≥ N ≥ 3 per group. Two independent experiments with similar results were performed; results are shown from one of them. Systemic leaves were harvested at 12 DPI for all assays. The same RNA extraction was used for all analyses for all samples.

### 3.3. TuMV enhances AGO2 and low-molecular weight forms of both AGO1 and AGO2 proteins

To investigate whether AGO1 or AGO2 protein levels were altered after TuMV infection, we performed Western blot analyses (**Fig. 5**). A visual inspection reveals that AGO1 remains at basal levels after TuMV infection (**Fig. 5a**), but AGO2 protein is greatly enhanced (**Fig. 5d**). Smaller forms recognized by antibodies against AGO1 and AGO2 proteins were also detected, which we termed AGO1* and AGO2*, respectively. Two independent experiments were performed and the results were combined using appropriate statistical tools (see Materials and Methods section). The results confirm and quantitatively supports the evidences previously seen after a naked eye inspection of the blots, namely, AGO1 protein levels were unchanged by TuMV infection (**Fig. 5b**) whereas AGO2 protein levels were greatly induced (**Fig. 5e**). Interestingly, both AGO1* and AGO2* products were consistently and highly induced by TuMV (**Fig. 5c,f**).

**Fig. 5.**
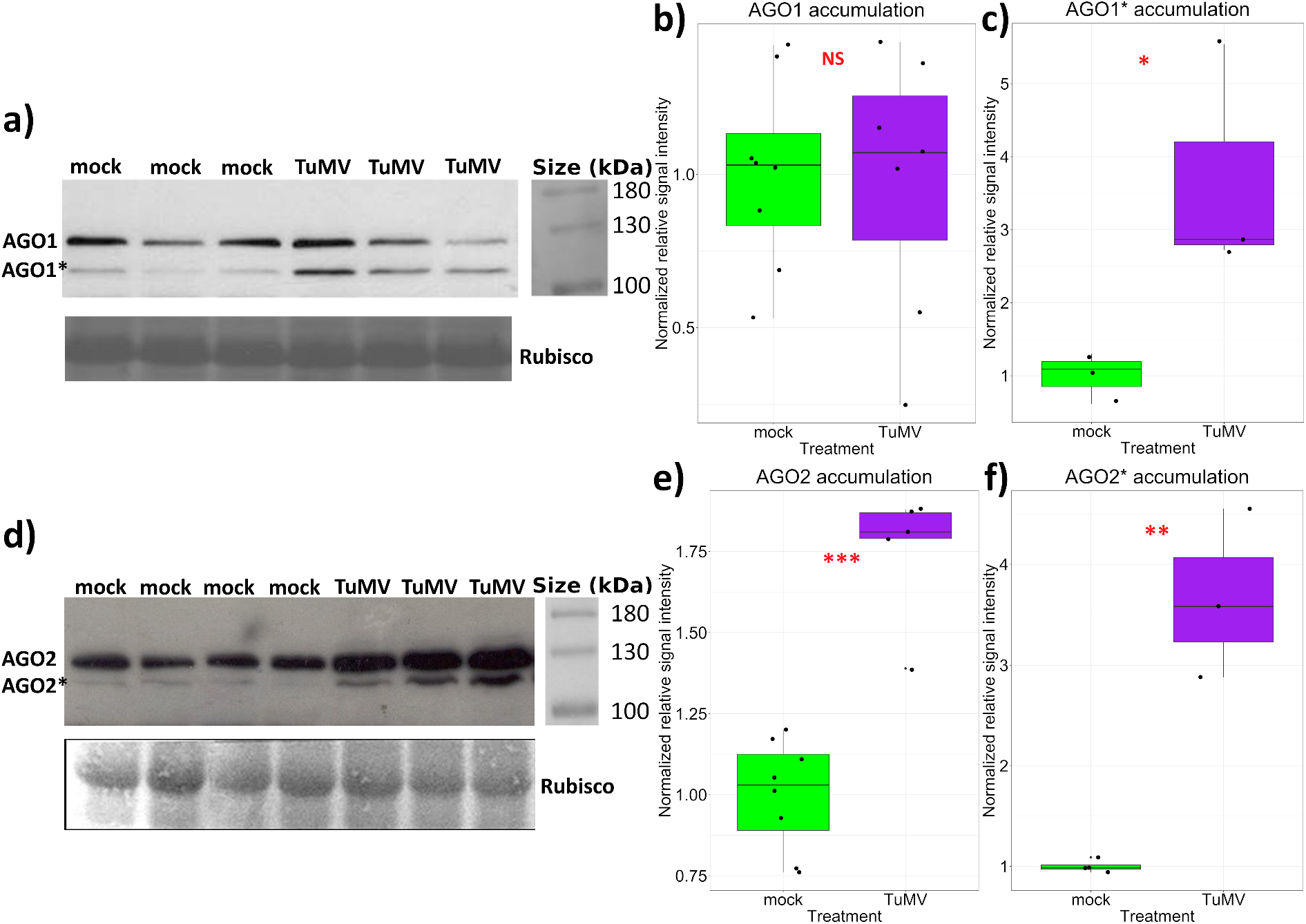
TuMV induces the accumulation of AGO2 protein and low-molecular weight forms of AGO1 and AGO2 proteins. Western blot analyses display accumulation of a) AGO1 and d) AGO2 proteins alongside their Rubisco loading control bands. Low-molecular weight forms (labeled AGO1* and AGO2*) are also shown. Two independent experiments that yielded similar results were performed and one representative experiment is depicted. Box-and-jitter plots of the combined results of two similar experiments display accumulation of b) AGO1, c) AGO1*, e) AGO2 and f) AGO2*.

## 4. Discussion

### 4.1. Simultaneous overaccumulation of miR168 and its cognate target AGO1 mRNA

Among plant microRNAs, the highly conserved miR168 is of paramount importance, as it directly regulates expression of the central miRNA effector, AGO1, ensuring proper functioning of the entire miRNA pathway (Vaucheret *et al*., 2004; Vaucheret *et al*., 2006). Through altering the endogenous miR168/AGO1 regulatory loop, plant viruses are thought to increase their fitness limiting the antiviral activity of AGO1 (Várallyay *et al*., 2010; Várallyay and Havelda, 2013; Harvey *et al*., 2011). As a result the endogenous miRNA pathway is altered as well (Yang and Li, 2018) and AGO2 becomes the main antiviral AGO protein (Harvey *et al*., 2011). Previous reports using TuMV-infected plants had showed overaccumulation of miR168 (Garcia-ruiz *et al*., 2015) and mRNA levels of AGO1/2/3 (Sánchez *et al*., 2015), separately. Here, we confirmed these findings both in long days (**Fig. 1, Fig. SF2, Fig. 2**) and short days (**Fig. 4**) conditions. Furthermore, we found that miR403 (which downregulates AGO2 mRNA) was also overaccumulated under TuMV (**Fig. 2, Fig. 4**).

TuMV suppressor of RNA silencing HC-Pro, is of utmost importance in determining interference in endogenous silencing AGO protein activity and concurrent symptoms development (Kasschau *et al*., 2003; Garcia-ruiz *et al*., 2015). Like other, unrelated viral suppressors, HC-Pro is capable by itself to trigger miR168 accumulation (Várallyay and Havelda, 2013).

Importantly, HC-Pro small RNA sequestering activity is not indiscriminate; while it massively sequesters viral-derived siRNAs (depleting AGO proteins from them and effectively impairing viral silencing), the proportion of endogenous miRNAs bounded by HC-Pro is very low (Garcia-ruiz *et al*., 2015). This is true also in the case of miR168 and miR403, which are overwhelmingly unbounded to HC-Pro (Garcia-ruiz *et al*., 2015). Therefore, rather than being stabilized by HC-Pro via physical interaction, we speculate that miR168 can be overaccumulated under TuMV infection via faster pre-miRNA processing (see below). Interestingly, TuMV-induced overaccumulation of mature miR168 does not lead to its increased loading into AGO1 neither (Garcia-ruiz *et al*., 2015). These findings using TuMV are in line with other reports that showed that mature miRNA accumulation for itself does not guarantee effective AGO1 loading/activity. MiR168 has a relatively very low rate of loading into RISC, remaining in high proportion unbounded as a cytoplasmic pool both in healthy plants and under viral infections, when it typically strongly over-accumulates (Dalmadi *et al*., 2019; Várallyay *et al*., 2010). Therefore, simultaneous over-accumulation of miR168 and its cognate target AGO1 mRNA (**Fig. 1, Fig. SF2, Fig. 4**), which appears counter-intuitive in principle, could be explained by ineffective RISC loading of surplus miR168.

The simultaneous overaccumulation of microRNAs and its cognate repressible mRNAs (termed “incoherent regulation”), has long been known for the regulatory loop miR168/AGO1 (Vaucheret *et al*., 2004) and was described under nutrient stress (Vidal *et al*., 2010; Jeong and Green, 2013), senescence (Thatcher *et al*., 2015) and viral infections (Várallyay *et al*., 2010; Zavallo *et al*., 2015). This phenomenon could arise from fine-tuned differentially spatio-temporally expression/localization of miRNAs and its targets (Válóczi *et al*., 2006). In fact, the question whether cells from different tissues respond differently to the same viral infection is an unanswered one and remains as a gap to fill in (Yang and Li, 2018). From a theoretical point of view, it was postulated that simultaneous activation of microRNAs and its targets is important to fine-tune the expression of the latter, which would otherwise underwent large stochastic expression (Osella *et al*., 2011).

### 4.2. TuMV triggers overaccumulation of mature miR168 and miR403 concomitantly with downregulation of its precursor forms

Transcriptional activity of pmiR168a was mostly unaffected by TuMV (**Fig. 1d, Fig. SF2d**). This result is in contrast to what (Várallyay and Havelda, 2013) found using other plant-virus interactions (though no potyvirus was used by these authors), and highlights the diversity of molecular responses following viral infections. However, our results are in line with those reported by other authors studying miRNA response to biotic and abiotic stresses. Interestingly, the enhanced accumulation of mature miR168 and miR403 (**Fig. 1a, Fig. SF2a, Fig. 2a, Fig. 4**) was paralleled by decreased accumulation of its corresponding precursor forms (**Fig. 1b-d, Fig. SF2b-d, Fig. 2b**). These results are similar to what (Dolata *et al*., 2016) found during salt stress when assessing with similar strategy the promoter activity of miR161 and miR173, which experienced unchanged transcriptional levels, but down-regulation of their pri-miRs forms concomitantly with an overaccumulation of their mature forms. These authors found that HYL1, a key player in miRNA biogenesis, was essential to trigger a differential pri-miRNA accumulation under salt stress. Similarly in rice, under RSV infection, (Zheng *et al*., 2017) found that the viral protein NS3, physically interacts with DRB1 (ortholog of Arabidopsis HYL1) aiding in pri-miRNA processing resulting in the simultaneous down-regulation of pri-miRNAs and enhanced accumulation of mature forms of several miRNAs associated with stress responses. These workers proposed that in this way, the virus can promote the processing of miRNAs whose targets are important genes in viral resistance or development. The fact that miR168a and b and miR403 are dependent upon HYL1 for its maturation (Szarzynska *et al*., 2009) raises the question whether the down-regulation of these pre-microRNAs under TuMV infections is a consequence of TuMV acceleration of miRNA processing via HYL1 and/or its molecular partners (Dong *et al*., 2008). Further studies should elucidate the role that HYL1 or other components of the plant microprocessor machinery could have in the alteration of the biogenesis of microRNAs 168 and 403, among others, during viral infections.

### 4.3. The role of ABA and SA in the silencing machinery following TuMV infection

ABA and SA increase under TuMV infection (Manacorda *et al*., 2021) and the regulatory systems studied here are sensitive to both hormones. For example, both miR168 and AGO1 are induced by ABA and ABA-responsive elements are present in the promoter sequences of these genes (Li *et al*., 2012; Alazem *et al*., 2019; Alazem *et al*., 2017). AGO1 showed a slight repression or no changes upon SA application ((Alazem *et al*., 2019); this study, **Fig. 3**). AGO2 is induced upon both SA or ABA treatments ((Alazem *et al*., 2019; Diao *et al*., 2019; Alazem *et al*., 2017); this study, **Fig. 3**). Therefore, TuMV-induced AGO1 and AGO2 mRNA accumulation (**Fig. 1e, Fig. SF2e, Fig. 2c, Fig. 4d, h**) is in good agreement with the expression profile of these genes under the effect of the hormones that TuMV greatly induces. AGO2 overexpression was detected earlier and this effect was more lasting than that of AGO1 (**Fig. 1e, Fig. 2c**). The importance of AGO2 as a key antiviral AGO protein in plants is being increasingly highlighted (Carbonell and Carrington, 2015; Ludman *et al*., 2017; Harvey *et al*., 2011). Interestingly, (Diao *et al*., 2019) applied a foliar MeSA spray to leaves founding that NbAGO2 showed a temporal response similar to what we found in **Fig. 3**. These authors also found that NbAGO2 mRNA accumulates upon TMV infection (which triggers SA accumulation) and is important in defense, confirming and expanding results obtained by (Ludman *et al*., 2017).

The balance between ABA and SA over miR168/AGO1 and miR403/ AGO2 systems could serve the plant to distinguish between abiotic and viral stresses: during abiotic stresses that trigger ABA, both AGO1 mRNA and miR168 are induced, generating a new equilibrium (Li *et al*., 2012), without SA induction that could trigger AGO2 strongly (Diao *et al*., 2019); this study, **Fig. 3**). During viral infections, often both ABA and SA play an antiviral role associated with the viral silencing machinery (Alazem *et al*., 2019). In this case, also ABA exerts its positive effect both on AGO1 mRNA and miR168 accumulation, but AGO1 is further limited by SA (Alazem *et al*., 2019). On the contrary, AGO2 is induced by both ABA and SA ((Alazem *et al*., 2019; Diao *et al*., 2019); this study, **Fig. 3**), escaping miR403 control at least in part. In this scenario, miR403, induced by TuMV (**Fig. 2a, Fig. 4e-g**), would not dispose of abundant AGO1 protein (**Fig. 5a, b**) to be loaded into, thus limiting its capacity to repress AGO2 mRNA and allowing AGO2 protein to overaccumulate (**Fig. 5d, e**) and exert its antiviral effect.

### 4.4. Differential effect of TuMV infection upon AGO1 and AGO2 protein accumulation

Regarding protein accumulation, we detected unchanged AGO1 levels upon infection (**Fig. 5a, b**) despite higher mRNA accumulation (**Fig. 1e, Fig. 4d**), a phenomenon reported by other workers after viral infections (Várallyay *et al*., 2010; Várallyay and Havelda, 2013). AGO2, on the contrary, was highly induced (**Fig. 5d, e**), again, an expected result based on what is known about AGO2 during viral infections (Harvey *et al*., 2011; Carbonell and Carrington, 2015; Ludman *et al*., 2017). The detection of lower molecular weight (MW) bands of AGO1/2 proteins had been reported previously (Qi and Mi, 2010; Harvey *et al*., 2011; Carbonell *et al*., 2012), but their nature remains unexplained. Here, we found that TuMV consistently increased the relative amount of these forms (**Fig. 5**) even in the case of AGO1, whose amount of complete protein is not significantly altered by TuMV. The identification of increased levels of low MW forms of AGO1/2 suggests that TuMV promotes the instability of both antiviral proteins upon infection. Further work should study the nature of these bands, their origin and the reasons for their increase during TuMV infection.

## 5. Conclusion

In sum, our work showed that TuMV alters both miR168/AGO1 and miR403/AGO2 regulatory systems at the RNA and protein levels. The simultaneous over-accumulation of both AGO1/2 mRNAs and their cognate respective repressors miR168 and miR403 (mature forms), adds another example to “incoherent regulation” under stress in plants. Similar transcriptional responses were detected both in long and short days conditions and no differences were detected between JPN1 and UK1 strains regarding these systems’ perturbation following viral infection. Early (4 DPI) response was weak, since only AGO2 mRNA experienced significant changes in accumulation. On the contrary, medium (10-12 DPI) and late (16 DPI) stages showed a strong response and very similar pattern of expression for the genes of these regulatory systems and highlighted that pre-miRNAs repression and mature miRNA over-accumulation is a long-lasting effect upon TuMV infection. Lack of AGO1 protein induction concomitantly with enhanced AGO2 over-accumulation stressed the prominent role of the latter as antiviral effector. Taking into account previous works that have focussed on abiotic/viral stress and the miRNA/siRNA pathways, this study highlights the importance of drawing links between the miRNA microprocessor machinery, the silencing machinery, and the impact that plant viruses could have on them due to hormonal disturbance.

## Captions for Supplementary Tables

**Supplementary Figure 1:** Arabidopsis plants growing in SD conditions showing characteristic TuMV symptoms at 12 DPI. (A) Plants were grown in trays to compare mock-inoculated plants (first two columns from the left) against TuMV-infected plants. Note the characteristic stunted and arrested growth, deepening of leaf serration and leaf twisting. Below, detailed photographs of either mock-inoculated (B) or TuMV-infected (C) Arabidopsis plants.

**Supplementary Figure 2:** MiR168/AGO1 system is similarly regulated by a TuMV strain with contrasting phenotypical outcomes. JPN1 strain of TuMV was used to replicate the experiment shown in **Fig.** 1. Identical conditions and **Fig.** captions apply as for **Fig.** 1.

**Supplementary Table 1:** MIQE Guidelines (*M*inimum *I*nformation for Publication of *Q*uantitative Real-Time PCR *E*xperiments) for mRNAs and miRNA Precursors.

**Supplementary Table 2:** MIQE Guidelines (*M*inimum *I*nformation for Publication of *Q*uantitative Real-Time PCR *E*xperiments) for microRNAs.

**Supplementary Table 3:** Oligonucleotide primers list.

## Author contributions

Conceptualization: Carlos A. Manacorda, María Rosa Marano and Sebastián Asurmendi; Methodology: Carlos A. Manacorda and Sabrina Tasselli; Formal analysis and investigation: Carlos A. Manacorda, Sebastián Asurmendi and Sabrina Tasselli; Writing - original draft preparation: Carlos A. Manacorda and Sebastián Asurmendi; Writing - review and editing: Carlos A. Manacorda, María Rosa Marano and Sebastián Asurmendi; Funding acquisition: Sebastián Asurmendi; Resources: Sebastián Asurmendi and María Rosa Marano; Supervision: Sebastián Asurmendi and María Rosa Marano.

## Acknowledgements

TuMV-UK1 strain (accession number AF169561) and JPN1 strain (accession number KM094174) were kind gifts of Dr. Flora Sánchez and Dr. Fernando Ponz. We also thanks Dr. Hervé Vaucheret for the kind gift of *pmiR168a:GUS* transgenic plants. Dr. Pablo Manavella and Dr. Nicolás Bologna provided technical advice in Northern blot and Western blot experiments, respectively, and technicians Agustín Montenegro, Matías Rodríguez and Ignacio Tévez aided with plant growth management. This work was supported by INTA PDi116 and by ANPCyT PICT2015 1532.

