## Supplementary figures and images for "TuMV infection alters miR168/AGO1 and miR403/AGO2 systems regulation in Arabidopsis"

### Supplemental Figure 1

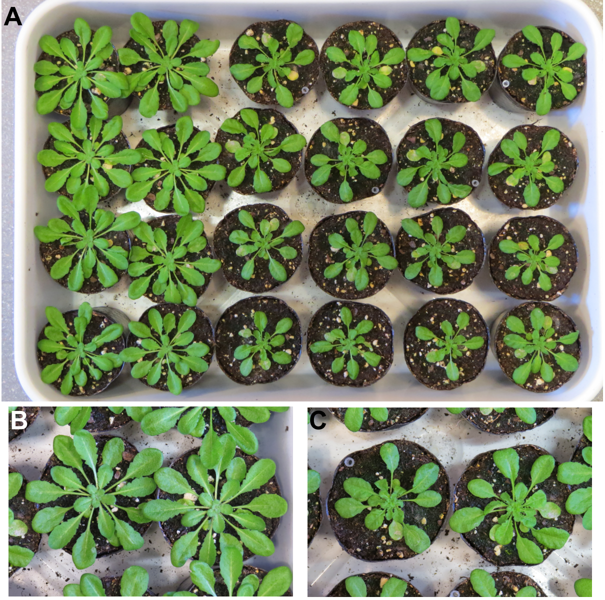

### Supplemental Figure 2

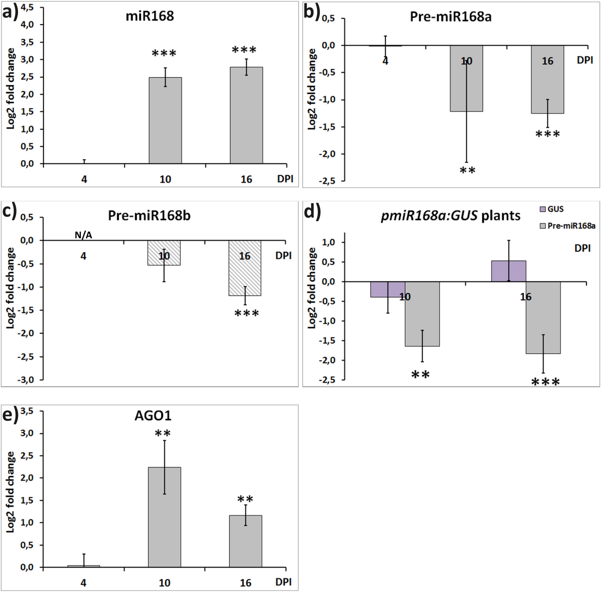
