## Supplemental Table 2 for "TuMV infection alters miR168/AGO1 and miR403/AGO2 systems regulation in Arabidopsis"

**Supplementary Table ST2:** Experimental conditions used for microRNA quantification by qRT-PCR assays based on MIQE requirements (Bustin et al., 2009; Bustin et al., 2010).

| **Experimental design** |  |
| --- | --- |
| Control groups | *A.thaliana* Col-0 buffer-inoculated |
| Treatment groups | *A.thaliana* Col-0 inoculated with either JPN1 or UK1 TuMV viruses |
| **Sample** |  |
| Type of sample | 11^th^ leaves (infection experiments in long day conditions) / Systemic rosette (leaves #8, #11 and #13, infection experiments in short day conditions). |
| Processing procedure | Liquid nitrogen homogenization |
| Sample frozen conditions | -80 ºC |
| Biological replicates | Leaves #11 used in infection experiments in long day conditions were pooled when necessary to obtain enough tissue for RNA extractions (4 DPI, 6≤n≤8; 10 DPI, n=2-3 for each pool). Each pool was treated as a separate biological replicate and qRT-PCR experiments were performed using 4≤n≤7 pools for each treatment and DPI / For systemic rosette leaves #8, #11 and #13, short day conditions plants, n= 3 plants were used per pool. |
| **RNA extraction (*)** |  |
| Procedure | Acid Phenol extraction |
| Reagents | TRIzol (Thermo Fisher Scientific) |
| Details of Dnase treatment | DNAse I Amp Grade (Thermo Fisher Scientific), 15 min at room temperature |
| Contamination assessment | < 3% |
| Nucleic acid quantification | Absorbance at 260 nm |
| Instrument and method | NanoDrop instrument |
| Purity( A260/ A 280) | > 2 (average) |
| Purity( A260/ A 230) | > 2 (average) |
| RNA integrity | Analyzed by agarose gel electrophoresis |
| **Reverse transcription (*)** |  |
| Complete reaction conditions | Based on the protocol developed by (Chen et al., 2005) and adapted by (Pant et al., 2008) to SYBR® Green reagents. |
| Amount of RNA and reaction volume | 100 ng of RNA, 20 µl |
| Priming oligonucleotide | Customized reverse-transcription primers for stem-loop amplification plus reverse primers for 4 selected reference genes. |
| Reverse transcriptase | SuperScript III Reverse Transcriptase (Thermo Fisher Scientific). |
| Temp and time | 30 min 16 ºC, (30 s 30 ºC, 30 s 42 ºC, 30 s 50 ºC) x 60 cycles, 5 min 85 ºC. |
| **qPCR protocol (*)** |  |
| Complete reaction conditions (**) | 5 min 95 ºC, (15 s 95 ºC, 1 min 60 ºC) x 45 cycles |
| Reaction volume and amount of cDNA | 2 µl of a 1/20 dilution of synthesized cDNA in a final volume of reaction of 20 µl. |
| Primers, Mg^2+^ and dNTPs concentration | 3 mM Mg^2+^, 200 nM primers, 0,2 mM dNTPs |
| Polymerase | Platinum® Taq DNA Polymerase (Thermo Fisher Scientific) |
| Buffer | 20 mM Tris-HCL (pH = 8.4), 50 mM KCl |
| Manufacturer of qPCR instrument | StepOnePlus, Thermo Fisher Scientific |
| **qPCR validation (*)** |  |
| Specificity | Analysed by agarose gel and Melting Curve parameters on each qPCR run and automatic MTP assessment by StepOnePlus software (v2.3). NTC assessment. |
| Method of PCR efficiency calculation | Mean PCR efficiency per amplicon calculated by LingRegPCR program (Ramakers et al., 2003). |
| **Data analysis (*)** |  |
| qPCR analysis program | LinRegPCR program |
| Method of Cq determination | LinRegPCR program |
| Outlier identification | LinRegPCR program |
| Justification of number and choice of reference genes | 4 reference genes tested (GAPC1 (NM_11283), TUB4 (NM_123801), UBQ5 (NM_116090) and EF1-α (NM_125432)) for stability using the Bestkeeper stability algorithm. UBQ5 was chosen as the most stable reference gene, as found in (Manacorda et al., 2013). |
| Description of normalization methods | (Pfaffl et al., 2002):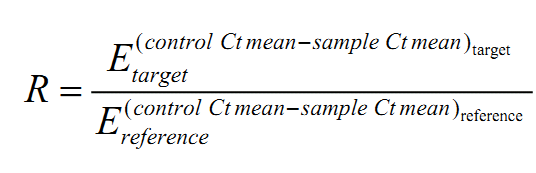 |
| Number of technical replicates | 2 |
| Statistical method | Permutation test |
| Software | fgStatistics software (Di Rienzo, 2009), (<http://sites.google.com/site/fgStatistics/>) |
| Repeatability (intraassay variation) C_q_ SD error | Between 0.08 and 0.36 depending on the assayed amplicon. |

(*) Details are given for long day conditions (leaves #11 of viral inoculation experiments). Similar protocols were followed for samples from short day conditions shown as Supplementary Data.

(**) Conditions are described for a generic qPCR assay. Particular annealing/extension temperatures could vary between amplicons. Specific conditions for each amplicon are available upon request.
